# Detection of endogenous MR1B protein in human 293F cells by immunoaffinity enrichment and two-dimensional western blotting

**DOI:** 10.64898/2026.10.17.759879

**Authors:** Hisateru Yamaguchi

**Affiliations:** Department of Management Nutrition, Aichi Gakusen University, 28 Kamikawanari, Hegoshi-cho, Okazaki, Aichi 444-8520, Japan; Institute for Comprehensive Medical Science, Fujita Health University, Toyoake, Aichi 470-1192, Japan

**Author notes:** **Corresponding Author:** Hisateru Yamaguchi, M.D., Ph.D. Department of Management Nutrition, Aichi Gakusen University, 28 Kamikawanari, Hegoshi-cho, Okazaki, Aichi 444-8520, Japan.

**Keywords:** MR1B, MR1, MAIT cells, splice variant, two-dimensional electrophoresis, immunoaffinity purification, HEK293

## Abstract

MR1B is a splice variant of the MHC class Ib molecule MR1 that lacks the α3 extracellular domain and has been reported to modulate MR1-mediated antigen presentation to mucosal-associated invariant T (MAIT) cells in transfected cells. Although MR1B transcripts are widely expressed, MR1B protein has been reported only in transfected overexpression systems; whether endogenous MR1B protein is present has remained unknown. Here we report the detection of endogenous MR1B protein. Immunoaffinity enrichment of 293F cell lysate (447 mg total protein) followed by two-dimensional electrophoresis and western blotting gave immunoreactive spots at the MR1A (∼40 kDa) and MR1B (∼27–30 kDa) reference coordinates established from recombinant cells, whose MR1 identity was confirmed by mass spectrometry. The spots were detected with two antisera raised against regions on opposite sides of the deleted segment and were absent with the corresponding pre-immune sera; detection with the C-terminal antiserum was obtained once. The 27–30 kDa spot lies on the basic side of the MR1A spot, the direction predicted for loss of the α3 domain and opposite to that predicted for a modeled, unmodified N-terminal fragment of similar mass. Mass spectrometry identified MR1 only in the recombinant reference spots, so the endogenous assignment rests on immunoreactivity and two-dimensional position. Endogenous MR1B protein is detected in 293F cells under non-overexpression conditions.

## Introduction

MR1 is a highly conserved, non-polymorphic MHC class Ib molecule encoded on chromosome 1q25, outside the classical MHC locus [1–3], which presents microbial riboflavin metabolites to mucosal-associated invariant T (MAIT) cells, an innate-like T cell population that recognises infected cells in an MR1-restricted manner [4, 5]. Besides the canonical full-length isoform MR1A, the locus encodes MR1B, a splice variant lacking the α3 extracellular domain [6].

MR1B transcripts are co-expressed with MR1A in diverse human tissues, often at comparable or higher levels, although the ratio varies among cell types and donors [7, 8]; both are likewise detected in commercial HEK293 cDNA by RT-PCR (data not shown), the transcript-level background to the present analysis. Functionally, co-expression of MR1B with MR1A in MR1-deficient cells suppressed MR1A-dependent antigen presentation to MAIT cells [8], whereas an earlier study reported that overexpressed MR1B activated MAIT cells in the presence of bacteria [9]; the function of MR1B thus remains unsettled, and both studies relied on transfected cells.

Despite this interest, MR1B protein has been reported only in transfected overexpression systems [7, 9]. Endogenous MR1 protein was not detected by western blotting of the cell lines and mouse tissues examined earlier [10], and in mouse a stabilising monoclonal antibody enabled its detection at the cell surface [11]. Since then endogenous MR1 has been followed in human cells with MR1-specific monoclonal antibodies and MR1-deficient controls by flow cytometry, imaging and T cell assays rather than by biochemical resolution of the isoforms, and because those antibodies recognise the α1-α2 platform shared by both, such studies do not distinguish MR1A from MR1B [12–14]. Endogenous MR1B protein itself has not been demonstrated.

Neither antiserum used here is MR1B-specific — both recognise regions retained by MR1A and MR1B — so isoform identity cannot be assigned from antibody reactivity alone, and one-dimensional blotting of an immunoaffinity eluate from parental 293F cells resolved no band attributable to endogenous MR1, a strong band of undetermined identity overlapping the MR1B region (Figure 1). We therefore combined immunoaffinity enrichment with two-dimensional electrophoresis, which separates the isoforms by their 10–13 kDa difference in apparent molecular weight along one axis and resolves co-migrating proteins by charge along the other, and assigned the resulting immunoreactive spots by positional concordance with recombinant references. The competition experiment of Figure 1 and the two-dimensional analysis of Figure 3 used separate preparations, on different immunoaffinity columns and different eluate pools (Materials and Methods).

**Figure 1.**
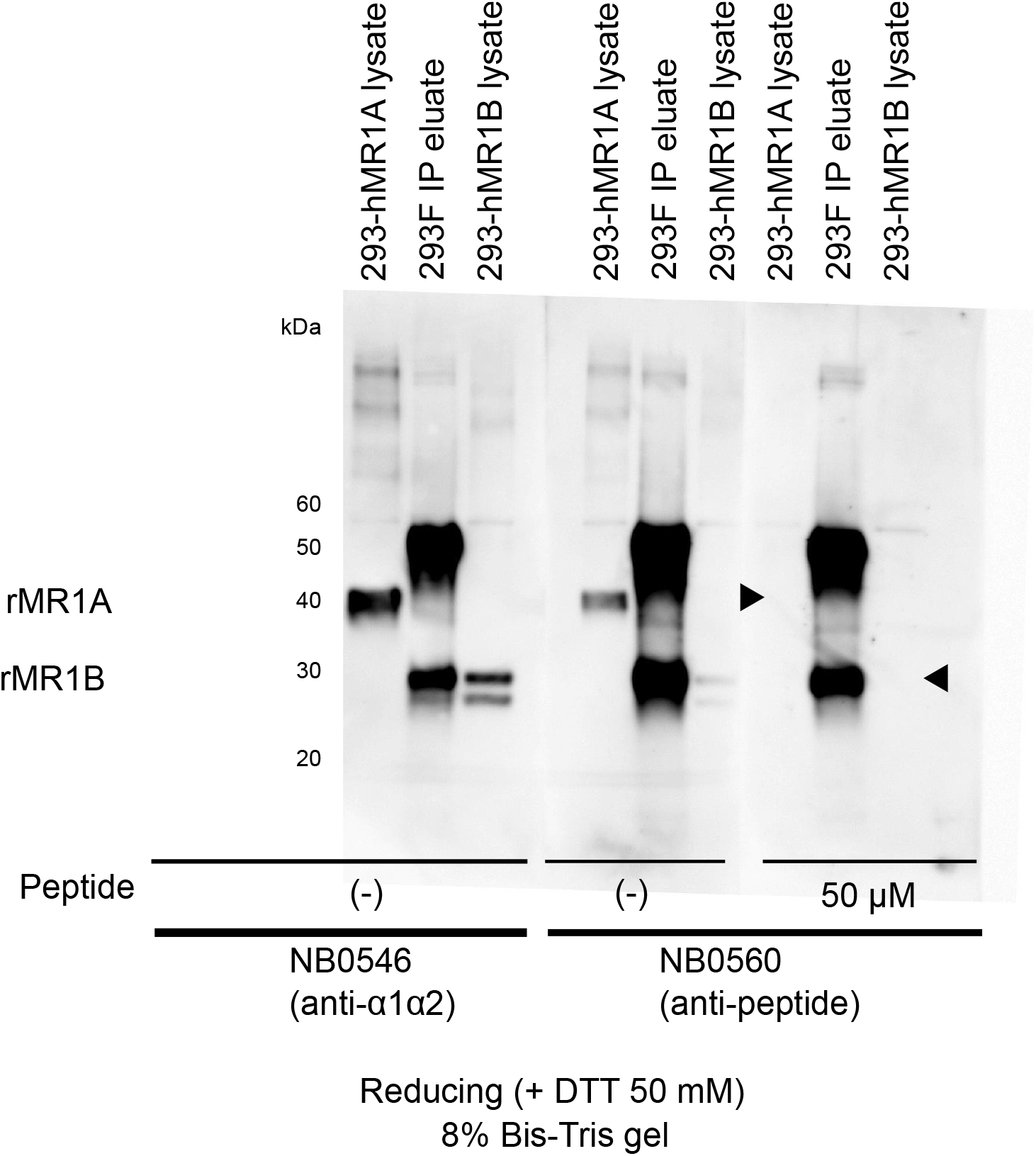
Peptide competition assay confirming NB0560 antiserum specificity. Lysates from 293-hMR1A and 293-hMR1B cells were resolved by one-dimensional SDS-PAGE (8% Bis-Tris, reducing) alongside a pooled immuno-precipitation eluate from parental 293F cells (lane order in each panel: 293-hMR1A lysate, 293F IP eluate, 293-hMR1B lysate). The membrane was cut into three strips, incubated with NB0546 (1:1,000; left panel) or NB0560 (1:500; centre and right panels), the latter without or with 50 μM cognate peptide (CRRPREQNGAIYLPTPDR), and imaged together. Marker and flow-through lanes were cropped from the displayed image, and molecular weight labels follow the marker lane of the same membrane. The abolition of the NB0560 signal was also seen in a separate experiment (data not shown). No band attributable to endogenous MR1 was resolved in the 293F lane with either antiserum; a prominent band there persisted with the peptide, overlapped the rMR1B region, and was not identified. Labels at the left mark the rMR1A (∼40 kDa) and rMR1B (∼27–30 kDa) levels; the arrowheads flanking the peptide-competed panel mark these levels beside the lanes in which each recombinant band appears without the peptide (rMR1A left, rMR1B right), where neither band remains.

## Results and Discussion

### Antiserum specificity and limits of one-dimensional detection

We first ran peptide competition assays on an immunoprecipitation eluate from parental 293F cells, with 293-hMR1A and 293-hMR1B lysates in the flanking lanes as molecular weight references (Figure 1). Pre-incubation of NB0560 with its cognate peptide abolished detection of both rMR1A (∼40 kDa) and rMR1B (∼27–30 kDa) in those lanes, showing that the NB0560 signal is peptide-specific, and its detection of rMR1B confirms that MR1B retains the C-terminal cytoplasmic region despite loss of α3. The same peptide did not affect NB0546 reactivity in an independent experiment (data not shown), consistent with NB0546 recognising the α1-α2 region. In the 293F eluate itself no band attributable to endogenous MR1 was resolved with either antiserum, and a prominent band that persisted with the competing peptide overlapped the rMR1B region; its identity was not determined. This background, with the lack of isoform-specific antisera, motivated the two-dimensional approach below.

### Two-dimensional reference coordinates for MR1A and MR1B

To establish reference positions for each isoform, eluates obtained from 293-hMR1A and 293-hMR1B lysates on an anti-MR1-C-term (NB0560) immunoaffinity column were resolved by two-dimensional electrophoresis and blotted with NB0546 (Figure 2a, b). The 293-hMR1A eluate gave multiple spots at approximately 40 kDa across a range of isoelectric points and the 293-hMR1B eluate multiple spots at approximately 27–30 kDa, consistent with deletion of the 92-residue α3 domain (predicted mass reduction approximately 10 kDa). Multiple spots per isoform rather than one discrete spot indicate charge heterogeneity; MR1 is N-glycosylated [7], but because the spots differ mainly in isoelectric point with little change in apparent molecular weight, the modification responsible cannot be identified from these data. The quantity used below is the direction of the charge difference between the isoforms, which agrees with calculation: for the mature chains of UniProt Q95460 isoform 1 (MR1A) and isoform 3 (MR1B), taken from precursor residue 23 with free termini and unmodified side chains, the EMBOSS pK set gives theoretical isoelectric points of 6.32 and 7.32, so loss of α3 is predicted to shift the protein approximately one unit towards the basic side. Absolute values depend on the pK set, the calculation omits glycosylation, and the non-linear strips do not allow an isoelectric point to be read off the gel, so only the position of a spot relative to the MR1A spot on the same gel is compared; because the gels differ in conditions (figure legends), positions were matched against the markers of each membrane to a width of a few kDa. The references appear as a train of several spots and the endogenous material as a smaller group; the terms spot pattern and spot are used accordingly.

**Figure 2.**
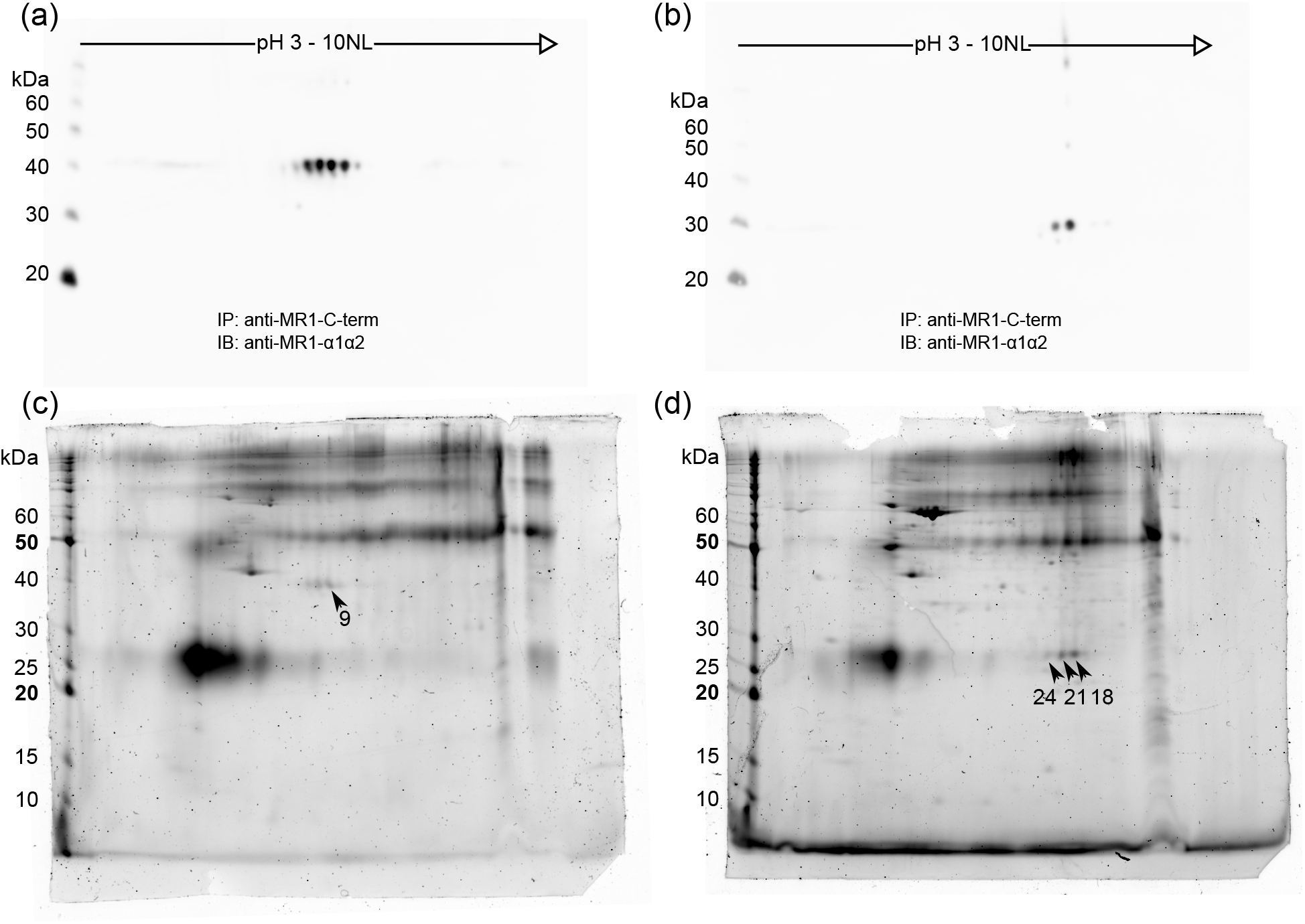
Two-dimensional electrophoresis of recombinant MR1A and MR1B. Eluates from anti-MR1-C-term (NB0560) immunoaffinity columns loaded with 293-hMR1A or 293-hMR1B lysate were resolved by two-dimensional electrophoresis (first dimension pH 3–10 NL except [c], pH 3–11 NL; second dimension 8% [a, b] or 10% [c, d] Bis-Tris). (a, b) 293-hMR1A and 293-hMR1B eluates immunoblotted with NB0546; multiple spots per isoform indicate charge heterogeneity (see text). rMR1A, ∼40 kDa; rMR1B, ∼27–30 kDa. (c, d) SYPRO Ruby-stained gels of the same two eluates, from separate immunoprecipitations, cropped from one fluorescence scan of the gels used for spot excision and rotated 90° clockwise to match (a, b); (c) was additionally rotated by approximately 1.8°. Numbered arrows, in (c) and (d) only, mark positions identified as MR1 by LC/MS/MS (numbers as in Table 1); in (d) the three arrows mark spots immediately above the 25 kDa marker (approximately 27 kDa), their labels placed below for legibility. Because the panels differ in gel conditions, positions were compared against the markers of each membrane to a width of a few kDa, not by overlaying gels. No spots were seen at the corresponding positions in the 293F gel (not shown). IP, immunoprecipitation; IB, immunoblot.

### Mass spectrometric confirmation of MR1 in recombinant reference spots

To verify that the recombinant material at the reference positions was MR1 protein, eluates from separate immunoprecipitations of 293-hMR1A and 293-hMR1B lysates were resolved on two-dimensional gels stained with SYPRO Ruby (Figure 2c, d), independently of the blots of Figure 2a, b, and spots at the expected molecular weights were excised for in-gel digestion and LC/MS/MS. MR1 was the top-ranked identification in four spots of these panels (Table 1), with Mascot protein scores up to 107, and in a spot from a separately prepared 293-hMR1B gel (not shown). A further such spot lay below the rMR1B reference position and was excluded from the positional comparison (spot numbers and scores in Table S1). No spots were visible at either reference position in the corresponding 293F gel processed in parallel. All identified peptides corresponded to sequences shared between MR1A and MR1B; peptides spanning the MR1B splice junction were not detected. Under the search settings recorded for these analyses, four junction-spanning peptides have unmodified neutral masses within 700–4000 Da, the window used for this comparison; all four terminate at Lys303 without extending into the following hydrophobic stretch (residues 304–323, canonical precursor numbering), whereas six others include the entire annotated transmembrane region (residues 303–323), two of them outside that window (Table S2). Their non-detection does not establish the absence of MR1B protein, and the factors that would determine whether they were observed — digestion, extraction, ionisation, precursor selection during data-dependent acquisition, and whether the junction sequence was in the searched database — were neither measured nor verified here.

**Table 1.**
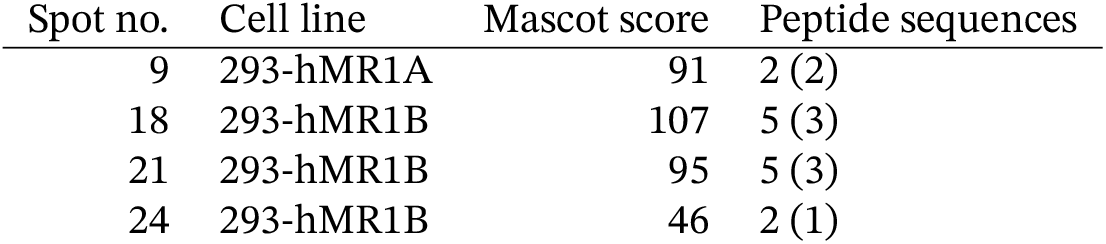
LC/MS/MS identification of MR1 protein in two-dimensional gel spots from recombinant cells. Spots excised from the panels of Figure 2c, d were digested in-gel with trypsin and analysed by nano-LC/MS/MS (Materials and Methods; search settings and full output in Table S1). Values are the Mascot protein score and the number of matched peptide sequences for the top-ranked MR1 hit, with the number significant at p < 0.05 in parentheses. All identified peptides are shared between MR1A and MR1B (see text).

### Immunoreactive spots at the MR1A and MR1B coordinates in 293F cells

We then applied the same approach to 293F lysate (447 mg total protein). Blotted with NB0546, the eluate gave two discrete spot patterns at the rMR1A (∼40 kDa) and rMR1B (∼27–30 kDa) reference coordinates (Figure 3c), and the independently raised NB0560 gave spot patterns at the same two positions (Figure 3d). As controls, aliquots of the same eluate were blotted with the corresponding pre-immune sera; neither produced a pattern at those positions (Figure 3a, b), although both showed non-specific background and IgG-related spots elsewhere, and because the same secondary antibody was used throughout, these panels control the contribution of immunoglobulin-derived species. The spots at the two positions appeared of broadly similar intensity on visual inspection, which is not used here to compare protein levels (see Data Limitations). With NB0546 the two-spot pattern was obtained in three independent immunoprecipitations (the large-scale preparation of Figure 3 and two smaller-scale experiments from 15 mg of lysate each, data not shown); the NB0560 blot was performed once, on the large-scale eluate.

**Figure 3.**
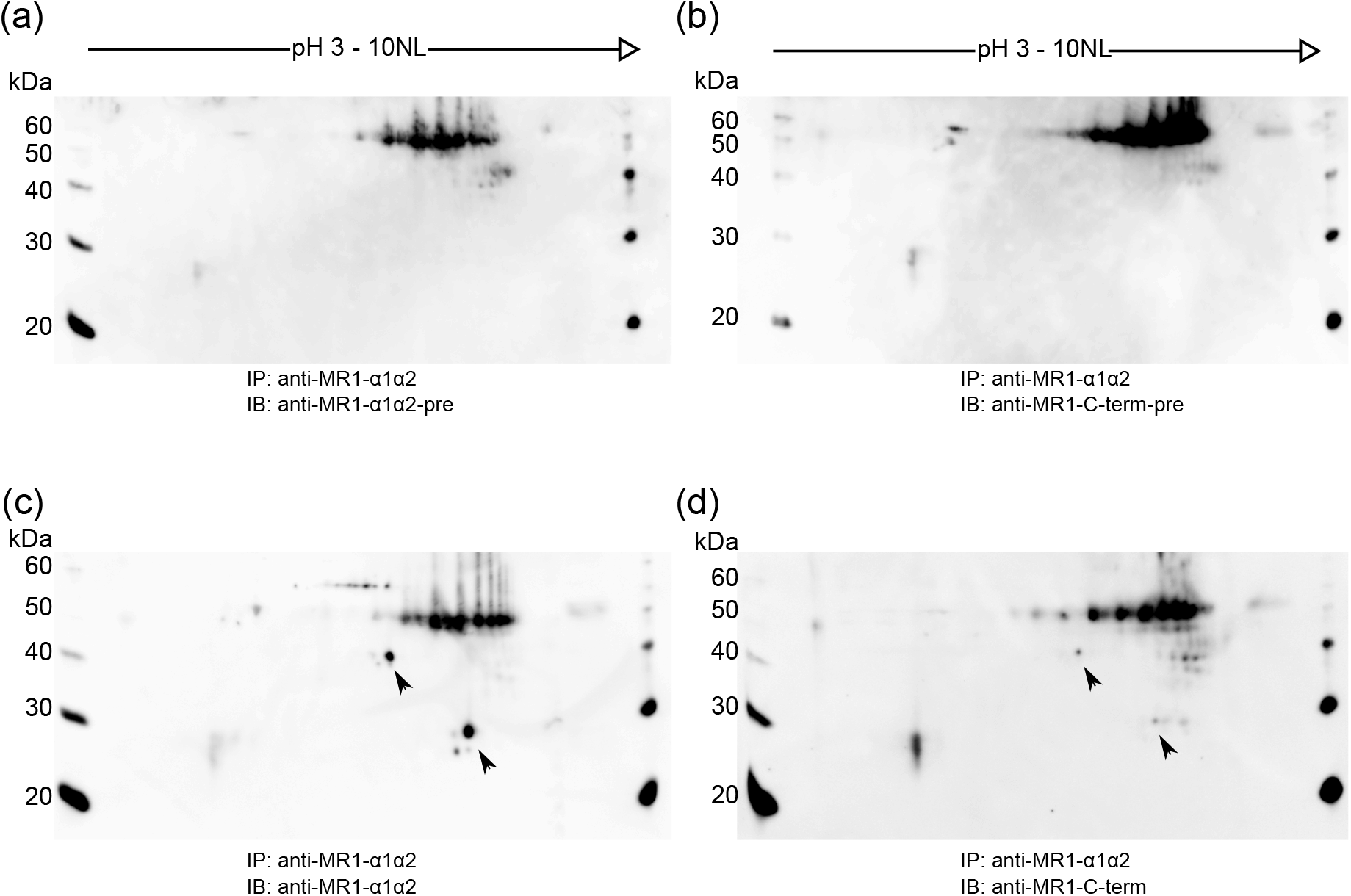
Two-dimensional western blotting of immunoprecipitation eluates from 293F cells. Eluate pooled from 447 mg of total lysate protein on the NB0546 column was resolved by two-dimensional electrophoresis (first dimension pH 3–10 NL; second dimension 10% Bis-Tris) and blotted. (a, b) Pre-immune sera corresponding to NB0546 and NB0560. (c) NB0546. (d) NB0560. Arrows mark the rMR1A (∼40 kDa) and rMR1B (∼27–30 kDa) reference coordinates of Figure 2a, b, matched against the markers of each membrane to a width of a few kDa; spots at the right edge of each panel are markers loaded at the basic end. All four panels are separate membranes prepared from aliquots of the same eluate, so reactivity with both antisera does not by itself show that the two recognised regions lie on one polypeptide chain. The pattern of (c) was obtained with NB0546 in three independent immunoprecipitations (the preparation shown and two smaller-scale experiments, data not shown); the NB0560 blot of (d) was performed once. IP, immunoprecipitation; IB, immunoblot; pre, pre-immune serum.

### Assignment of the 27-30 kDa species to MR1B

The immunoreactive species at the 27–30 kDa coordinate satisfies three conditions at once: its apparent mass is reduced by approximately 10 kDa relative to MR1A, it is labelled by two antisera raised against regions on opposite sides of the deleted segment, and it lies on the basic side of the MR1A spot. Taken together, these are the properties expected of an internal deletion of the α3 domain, and the alternative explanations available from the present data can be compared against them as follows. If the NB0560-reactive species is an MR1-derived polypeptide recognized through its native C-terminal sequence, a simple C-terminal truncation accounting for the mass loss would remove that sequence and would therefore be inconsistent with this interpretation. A truncation at the other end cannot be addressed through the epitopes, because NB0546 is polyclonal and the region of its immunogen (mature residues 1–179) that it recognises has not been mapped; it can, however, be compared with the predicted charge. Modelled as unmodified mature chains, fragments spanning precursor residues 108–341 and 110–341 have computed masses of 26.71 and 26.48 kDa, close to that of MR1B (26.53 kDa), but theoretical isoelectric points of 4.96 and 4.84 — approximately 1.4 units acidic of the MR1A mature chain, where MR1B lies approximately 1.0 unit basic of it. The observed species lies basic of MR1A, so its position is opposite in sign to the prediction for these modelled fragments; this applies to the fragments as specified and does not exclude N-terminally truncated species in general. Among the other annotated isoforms of UniProt Q95460, MR1C lacks the native C-terminal immunogen sequence altogether, so specific recognition through that sequence is not predicted for it, although cross-reactivity has not been excluded experimentally; isoform 2 (31.84 kDa) lies above the observed window and isoform 5 (22.75 kDa) below it, and their calculated charge differences from MR1A (+0.48 and +0.28 unit) are smaller than that of MR1B (+1.00 unit). Computed masses of unmodified chains are not the same quantity as apparent migration masses of glycosylated proteins, and these comparisons are made in that light.

Why endogenous MR1B has been difficult to detect is apparent from the same data. In overexpression systems the abundance of the protein and the known identity of the transfected construct make both detection and isoform assignment straightforward, whereas here no band attributable to endogenous MR1 could be resolved in the one-dimensional blot of the 293F eluate, in which a strong peptide-insensitive band of undetermined identity overlapped the MR1B region (Figure 1, centre lane); and neither antiserum is isoform-specific, so isoform identity depends on positional information. Immunoaffinity enrichment combined with two-dimensional electrophoresis addresses both points, and previous reports of MR1B protein relied on transfected overexpression systems [7, 9], which the present observations extend to non-overexpression conditions in 293F cells.

Four limits of this assignment should be stated with it. Figure 3c and Figure 3d are separate membranes prepared from the same eluate, so reactivity with both antisera is not direct evidence that the two recognised regions lie on a single polypeptide chain; nothing in these data demonstrates the splice junction itself; the pre-immune controls do not exclude reactivity that arose after immunisation; and the peptide competition of Figure 1 was performed on the recombinant lanes, not on the candidate spots of Figure 3. This is, to our knowledge, the first detection of endogenous MR1B protein, in human 293F cells without overexpression.

### Data Limitations and Perspectives

Five limitations should be stated. First, the evidence is immunochemical and positional, not peptide-level: junction-spanning peptides were not detected, and MR1 was identified by mass spectrometry only in the recombinant reference spots. LC/MS/MS of spots from SYPRO Ruby-stained gels of parental 293F eluates in two earlier experiments identified immunoglobulin-derived peptides but not MR1; those preparations differ from the one of Figure 3 and do not establish the composition of its spots. Second, relative abundance was not evaluated: chemiluminescent signal is not linear and recovery and antiserum reactivity were not calibrated, so these intensities do not support a comparison of protein levels, which would require fluorescence detection or quantitative proteomics. Third, the antisera are polyclonal homemade reagents whose sensitivity at endogenous levels cannot be quantified, although both detect recombinant MR1A and MR1B on blots (Figures 1 and 2a, b) and NB0546 detects MR1B at the cell surface by flow cytometry [7]. Fourth, the observations come from a single HEK293-derived cell line. Fifth, no samples, antisera, cell lines or constructs from this study remain for further experiments.

Direct confirmation will require MR1B-specific antibodies, for example against the splice-junction region, or targeted mass spectrometry of the junction peptides. Of the four candidates of Table S2 the fully cleaved peptide TESETIPLVMK (monoisotopic 1246.65 Da; 2+ m/z 624.33) is the first choice, and an assay would have to specify the modification states monitored, such as oxidised Met. The four are nested sequences sharing the junction, Met302 and the C-terminal Lys303 and differing only in missed cleavages on their N-terminal side, so they are neither four independent lines of evidence nor four detection opportunities of equal efficiency. They are specific to MR1B among the annotated isoforms of UniProt Q95460, but their proteome-wide specificity, recovery, retention, junction-diagnostic fragment ions and performance in real samples are untested: the list proposes candidates, not a validated assay. Confirmation in primary cells and other cell types, and in cells lacking MR1, remains to be done.

## Materials and Methods

Buffer compositions, the preparation of the immunoaffinity column, the isoelectric focusing and SDS-PAGE programmes, blotting and development, instrument settings and the in-gel digestion procedure are given in Supplementary Methods; this section states what the observations reported here rest on.

### Cell lines and antibodies

FreeStyle 293-F cells (293F; Thermo Fisher Scientific; RRID: CVCL_D603), a suspension-adapted derivative of HEK293 cells, were the source of endogenous MR1 protein, and stable 293F transfectants expressing untagged human MR1A or MR1B (293-MR1A and 293-MR1B in [7]; here 293-hMR1A and 293-hMR1B) served as references, their MR1 proteins being called rMR1A and rMR1B. Two rabbit polyclonal antisera were used throughout [7]. Anti-MR1-α1α2 antiserum (NB0546) was raised against a recombinant protein encompassing the MR1 α1–α2 domains (mature residues 1–179). Anti-MR1-C-term antiserum (NB0560) was raised against the 18-residue synthetic peptide CRRPREQNGAIYLPTPDR, in which the 17 C-terminal residues RRPREQNGAIYLPTPDR correspond to the cytoplasmic sequence of MR1 (precursor residues 325–341) and the N-terminal cysteine is not part of the native sequence. Both regions are retained by MR1A and MR1B. One goat anti-rabbit IgG–HRP conjugate was used as secondary antibody throughout.

### Immunoprecipitation

293F cells were lysed in 1% CHAPS buffer. For Figure 3, 447 mg of total lysate protein was processed in five consecutive batches on a column of IgG purified from NB0546 and the concentrated eluates were pooled, a 100 μl aliquot (approximately 36 mg of lysate equivalent) being used per two-dimensional analysis; two earlier, smaller-scale experiments processed 15 mg of lysate each on the same column, which was used only for parental 293F lysate. For Figure 1, six stored 293F eluate fractions were pooled and concentrated. The recombinant references were prepared on NB0560 columns from 2 mg of 293-hMR1A or 293-hMR1B lysate for Figure 2a, b, and from 3–4 mg on a cross-linked column as described [7] for Figure 2c, d.

### Electrophoresis, western blotting and mass spectrometry

Samples were resolved by isoelectric focusing followed by SDS-PAGE, with molecular weight markers run on every gel, and blots were developed by enhanced chemiluminescence; for the controls the respective pre-immune serum replaced the primary antiserum at the same dilution. For the peptide competition assay, NB0560 or NB0546 was pre-incubated with 50 μM of the C-terminal peptide for 1 h at room temperature before use as primary antibody. Gels for mass spectrometry were stained with SYPRO Ruby, and excised spots were digested in-gel with trypsin and analysed by nano-LC/MS/MS on an ion-trap instrument, essentially as described [15]. For the recombinant reference spots (Table 1), Mascot MS/MS ions searches were run against NCBInr version 20101020 (taxonomy Homo sapiens) with trypsin and up to three missed cleavages; for the two earlier searches of parental 293F eluates, see Data Limitations.

### RT-PCR

RT-PCR of commercial HEK293 cDNA (Human Cell Line MTC panel, Clontech) used the primers and conditions described previously [7], with 40 cycles for MR1 and 22 for GAPDH.

## Acknowledgments

The author thanks Akihiko Takasaki (then at Institute for Comprehensive Medical Science, Fujita Health University, Toyoake, Japan) for performing the LC/MS/MS analysis.

## Author Contributions

H.Y.: Conceptualization, Data curation, Investigation, Methodology, Visualization, Writing – original draft, Writing – review and editing.

## Data Availability

The experimental data supporting the findings of this study are available from the corresponding author upon reasonable request. Supplementary material comprises Supplementary Methods, the Mascot output and search settings for the spots of Table 1 (Table S1), the in silico junction-spanning peptides of MR1B (Table S2), and uncropped membranes with exposure series for Figures 1–3 (Figures S1–S3).

## Declaration of Generative AI Use

The author used Claude Code (Anthropic) for text drafting, structural organisation and reference management, and Codex (OpenAI) and Antigravity (Google) for independent verification of the text and figures. The author reviewed and edited all content, made all scientific and interpretive decisions, and takes full responsibility for the publication.

## References

[1] K. Hashimoto, M. Hirai, Y. Kurosawa, A gene outside the human MHC related to classical HLA class I genes, Science 269 (1995) 693–695. 10.1126/science.7624800.

[2] H. Yamaguchi, M. Hirai, Y. Kurosawa, K. Hashimoto, A highly conserved major histocompatibility complex class I-related gene in mammals, Biochem. Biophys. Res. Commun. 238 (1997) 697–702. 10.1006/bbrc.1997.7379.

[3] H. Yamaguchi, Y. Kurosawa, K. Hashimoto, Expanded genomic organization of conserved mammalian MHC class I-related genes, human MR1 and its murine ortholog, Biochem. Biophys. Res. Commun. 250 (1998) 558–564. 10.1006/bbrc.1998.9353.

[4] H.E.G. McWilliam, J.A. Villadangos, MR1 antigen presentation to MAIT cells and other MR1-restricted T cells, Nat. Rev. Immunol. 24 (2024) 178–192. 10.1038/s41577-023-00934-1.

[5] D.I. Godfrey, H.-F. Koay, J. McCluskey, N.A. Gherardin, The biology and functional importance of MAIT cells, Nat. Immunol. 20 (2019) 1110–1128. 10.1038/s41590-019-0444-8.

[6] P. Riegert, V. Wanner, S. Bahram, Genomics, isoforms, expression, and phylogeny of the MHC class I-related MR1 gene, J. Immunol. 161 (1998) 4066–4077. 10.4049/jimmunol.161.8.4066.

[7] H. Yamaguchi, K. Tsukamoto, K. Hashimoto, Cell surface expression of MR1B, a splice variant of the MHC class I-related molecule MR1, revealed with antibodies, Biochem. Biophys. Res. Commun. 443 (2014) 422–427. 10.1016/j.bbrc.2013.11.096.

[8] G.A. Narayanan, A. Nellore, J. Tran, et al., Alternative splicing of MR1 regulates antigen presentation to MAIT cells, Sci. Rep. 10 (2020) 15429. 10.1038/s41598-020-72394-9.

[9] J. Lion, V. Debuysscher, A. Wlodarczyk, et al., MR1B, a natural spliced isoform of the MHC-related 1 protein, is expressed as homodimers at the cell surface and activates MAIT cells, Eur. J. Immunol. 43 (2013) 1363–1373. 10.1002/eji.201242461.

[10] H. Yamaguchi, K. Hashimoto, Association of MR1 protein, an MHC class I-related molecule, with β2-microglobulin, Biochem. Biophys. Res. Commun. 290 (2002) 722–729. 10.1006/bbrc.2001.6277.

[11] W.-J. Chua, S. Kim, N. Myers, et al., Endogenous MHC-related protein 1 is transiently expressed on the plasma membrane in a conformation that activates mucosal-associated invariant T cells, J. Immunol. 186 (2011) 4744–4750. 10.4049/jimmunol.1003254.

[12] H.E.G. McWilliam, S.B.G. Eckle, A. Theodossis, et al., The intracellular pathway for the presentation of vitamin B-related antigens by the antigen-presenting molecule MR1, Nat. Immunol. 17 (2016) 531–537. 10.1038/ni.3416.

[13] B. Laugel, A. Lloyd, E.W. Meermeier, et al., Engineering of Isogenic Cells Deficient for MR1 with a CRISPR/Cas9 Lentiviral System: Tools To Study Microbial Antigen Processing and Presentation to Human MR1-Restricted T Cells, J. Immunol. 197 (2016) 971–982. 10.4049/jimmunol.1501402.

[14] M. Salio, W. Awad, N. Veerapen, et al., Ligand-dependent downregulation of MR1 cell surface expression, Proc. Natl. Acad. Sci. USA 117 (2020) 10465–10475. 10.1073/pnas.2003136117.

[15] G. Kurosawa, M. Sumitomo, Y. Akahori, et al., Methods for comprehensive identification of membrane proteins recognized by a large number of monoclonal antibodies, J. Immunol. Methods 351 (2009) 1–12. 10.1016/j.jim.2009.09.003.

