## Supplementary Material for "Detection of endogenous MR1B protein in human 293F cells by immunoaffinity enrichment and two-dimensional western blotting"

Hisateru Yamaguchi[a,b]

#### Supplementary Methods

Conditions are given for the experiments shown in Figures 1–3 and Table 1, which are identified here by date.

##### Cells and lysis

FreeStyle 293-F cells and the stable 293-hMR1A and 293-hMR1B transfectants were maintained as described [1]. Cells were lysed in 1% CHAPS lysis buffer: 1% (w/v) CHAPS, 10 mM Tris-HCl (pH 7.8), 150 mM NaCl, 1 mM EDTA. Total protein was measured with the Micro BCA Protein Assay Kit (Thermo Fisher Scientific). For the recombinant eluates of Figure 2c, d the lysis buffer was TNF buffer: 10 mM Tris-HCl (pH 7.8), 150 mM NaCl, 1 mM EDTA, 1% NP-40.

##### Immunoaffinity columns

Three columns were used, one for each group of preparations below.

The NB0546 column (Figure 3, and the two smaller-scale replicates) was prepared on 2014-06-03 and used only for parental 293F lysate. IgG was purified from 2 ml of antiserum NB0546 on Protein G Sepharose 4 Fast Flow (GE Healthcare) and dialysed against 0.2 M $NaHCO_3$, 0.5 M NaCl (pH 8.3), giving 8.4 mg of IgG (2.4 mg/ml, 3.5 ml). The IgG was coupled to 1.5 ml of NHS-activated Sepharose 4 Fast Flow (GE Healthcare, catalogue no. 17-0906-01) by rotation overnight at 4 °C, blocked with 0.1 M Tris-HCl (pH 8.5), 0.5 M NaCl for a few hours at room temperature, washed alternately with that buffer and 0.1 M sodium acetate (pH 4.0), 0.5 M NaCl, and stored in PBS containing 0.1% $NaN_3$ at 4 °C.

The NB0560 column was prepared on 2014-02-21 from approximately 8 mg of Protein G-purified NB0560 IgG on 1 ml of NHS-activated Sepharose by the same procedure.

The recombinant reference eluates of Figure 2c, d were prepared on NB0560 antiserum bound to Protein G Sepharose and cross-linked with dimethyl pimelimidate, and were eluted with 0.1% trifluoroacetic acid, as described [1].

##### Immunoprecipitation and concentration

Lysate was rotated with the column overnight at 4 °C and the flow-through collected. The column was then washed with 1% CHAPS lysis buffer (40 ml in the large-scale runs) and eluted with 0.1 M glycine-HCl (pH

2.5), approximately 8 ml being collected per run. The column was neutralised with PBS and re-used. Eluates were concentrated on Amicon Ultra-4 units (Millipore, Ultracel 3 kDa cut-off, catalogue no. UFC800396) at 7,500 × g, each batch separately.

For Figure 3, 447 mg of 293F lysate protein was processed in five consecutive batches on the NB0546 column, each eluate concentrated to approximately 250 µl and the five pooled to 1.25 ml. A 100 µl aliquot of this pool, containing 44 µg of protein before clean-up and corresponding to approximately 36 mg of lysate, was used for each two-dimensional analysis. Each aliquot was taken through the 2-D Clean-Up Kit (GE Healthcare), procedure A for 1–100 µl samples containing 1–100 µg of protein, and the pellet was resuspended in 130 µl of rehydration solution. The two smaller-scale replicates processed 15 mg of lysate each on the same column.

For Figure 1, six stored 293F eluate fractions (recorded as 140226, 140327, 140408, 140421, 140430 and 140512) were concentrated together to approximately 150 µl on 2014-05-13, five of them from immunoprecipitations on the NB0560 column eluted with 0.1 M glycine-HCl (pH 2.5). For the blot, 10 µl per lane was loaded from a 43 µl sample prepared from 30 µl of the pool.

For Figure 2a, b, 1 ml of 293-hMR1A or 293-hMR1B lysate at 2 mg/ml was processed on the NB0560 column and eluted in 200 µl, of which 50 µl was loaded per gel. For Figure 2c, d, 4 mg (Figure 2c) and 3 mg (Figure 2d) of the corresponding lysates in TNF buffer were processed on the cross-linked column, eluted with 0.1% trifluoroacetic acid and lyophilised.

#### Two-dimensional electrophoresis

The rehydration stock solution was 2 M thiourea, 7 M urea, 4% (w/v) CHAPS in water. The working rehydration solution was prepared from this stock with the IPG buffer matching the strip, DTT and bromophenol blue, for example 150 µl of stock, 10 µl of 16% (w/v) DTT, 1.3 µl of IPG buffer and 2 µl of bromophenol blue.

The first dimension used Immobiline DryStrip gels, 7 cm, pH 3–10 NL or pH 3–11 NL (GE Healthcare, catalogue no. 17-6003-73) with the matching IPG Buffer, on an IPGphor unit at 20 °C and 50 µA per strip. The programme of the 2014 runs was: rehydration at 0 V (2 h in the holder at 20 °C and a further 3 h for the 2014-08 and 2014-11 runs; 2 h for the 2014-05 run), 30 V step-and-hold for 12 h, 300 V step-and-hold to 200 Vh, gradient to 1,000 V over 300 Vh, gradient to 5,000 V over 4,500 Vh, 5,000 V step-and-hold to 3,000 Vh, and 300 V step-and-hold for up to 12 h. The 2010 runs used the same steps except that the 300 V step was run to 750 Vh and the 5,000 V hold to 2,000 Vh, with no additional rehydration step. The totals recorded at the end of focusing were 9,215 Vh (2014-05-19, Figure 2a, b), 8,533 Vh (2014-08-13, Figure 3c), 8,812 Vh (2014-11-10, Figure 3d), 9,358 Vh (2014-11-12, Figure 3a, b), 8,807 Vh (2010-10-04, Figure 2c) and 8,721 Vh (2010-10-13, Figure 2d).

Focused strips were equilibrated for 15 min in equilibration buffer containing DTT at 10 mg/ml and then for 15 min in the same buffer containing iodoacetamide at 25 mg/ml. The equilibration buffer was 50 mM Tris-HCl (pH 7.5), 6 M urea, 30% glycerol, 2% (w/v) SDS, 0.002% bromophenol blue; the pH was set near neutral to match the Bis-Tris gel system. Strips were sealed onto the second-dimension gel with 0.5% (w/v) low-melting agarose in MOPS running buffer.

The second dimension was run in MOPS buffer at 200 V in an XCell SureLock Mini-Cell (Invitrogen) on Bis-Tris gels: 8% for Figure 2a, b, for the one-dimensional gel of Figure 1 and for the two smaller-scale replicates, and 10% for Figures 2c, d and 3. Markers were loaded alongside the strip at the acidic (+) end of the second-dimension gel: MagicMark XP Western Protein Standard (Life Technologies) for gels to be transferred to PVDF membrane and BenchMark Unstained Protein Ladder (Life Technologies, catalogue no. 10747-012) for gels designated for mass spectrometry. The recorded amounts are 0.3 µl at the acidic and 0.2 µl at the basic end for MagicMark XP, and 2 µl and 1 µl respectively for BenchMark. For the Figure 3 gels, MagicMark XP was applied at both ends of the second-dimension gel, and the spots at the right-hand edge of those panels are this marker.

For the one-dimensional gel of Figure 1, lysates of 293-hMR1A and 293-hMR1B were loaded at 0.5 μg per lane and the 293F eluate at 10 μl per lane, in LDS sample buffer with DTT, heated at 70 °C for 10 min.

#### Western blotting and development

Gels were transferred to PVDF membrane at 30 V for 60 min in an XCell II Blot Module (Invitrogen), rinsed in PBS for 10 min and blocked in 2% (w/v) skim milk (Wako), 0.1% Tween-20, PBS: 1 h for the blot of Figure 1 and overnight for the two-dimensional blots. Primary antisera were diluted in PBS with 0.1% Tween-20: NB0546 at 1:1,000 and NB0560 at 1:500 for Figure 1; NB0546 at 1:1,000 for Figure 2a, b and Figure 3c; NB0560 at 1:1,000 for Figure 3d; and the pre-immune sera corresponding to NB0546 and NB0560 at 1:1,000 for the controls of Figure 3a, b. The secondary antibody was a goat anti-rabbit IgG (H+L)–HRP conjugate (MBL, catalogue no. 458) at 1:10,000 for 30 min, used for every panel of Figures 1–3. Blots were developed with Pierce Western Blotting Substrate Plus (Thermo Fisher Scientific, product no. NCI32132JP) and imaged on an ImageQuant LAS 4000. Each membrane was imaged at 10, 30, 90 and 270 s and as a 400 s cumulative image; these are shown in Figures S1–S3.

#### Peptide competition assay

The competing peptide was the NB0560 immunogen CRRPREQNGAIYLPTPDR, added to the diluted primary antiserum to 50 μM from a 10 mg/ml stock (for example 300 μl of diluted serum plus 3.21 μl of stock) and pre-incubated for 60 min at room temperature before the membrane was probed. In the independent experiment of 2014-03-28 both antisera were tested with and without the peptide in parallel; this is the experiment referred to in the text as showing that NB0546 is unaffected by it.

#### Gel staining, imaging and spot excision

Gels designated for mass spectrometry were washed in water for 10 min, fixed twice for 30 min in 50% methanol, 7% acetic acid, washed in water for 10 min, stained with SYPRO Ruby overnight, destained for 30 min in 10% methanol, 7% acetic acid and washed in water for 10 min. Gels were scanned on a Typhoon imager (GE Healthcare). For excision, six gels were laid out together, fixed around the edges with 1% agarose and provided with reference markers at both ends, and scanned at a 100 μm pixel size; a pick list was generated in DeCyder and the spots were excised with an Ettan Spot Picker into 100 μl of water in eight-tube PCR strips, after which the gels were rescanned to confirm which positions had been taken. The spot numbers of Table 1 and Table S1 are the sequential positions of this pick list.

For the two endogenous analyses of 2014 (2014-06 and 2014-07), spots were excised manually from SYPRO Ruby-stained gels of 293F eluates rather than robotically.

#### In-gel digestion

Excised gel pieces were destained three times for 10 min at room temperature in 150 μl of 25 mM ammonium bicarbonate, 50% acetonitrile, dehydrated twice for 10 min in 100 μl of acetonitrile and air-dried for 20–60 min. Reduction and alkylation were omitted at this step; the carbamidomethylation of cysteine included in the search settings comes from the iodoacetamide of the strip equilibration. Digestion used trypsin at 20 μg/ml in 50 mM ammonium bicarbonate, enough to cover the gel piece (5 μl in 2010 and 15 μl in 2014), 30 min on ice followed by overnight incubation at 37 °C in capped tubes. Peptides were extracted with 10 μl of 1% trifluoroacetic acid for 10 min followed by 25–30 μl of acetonitrile for 10 min, in three rounds, and concentrated in a vacuum centrifuge; the volume was then adjusted to 40 μl with water and 4 μl of a stock of 20% acetonitrile, 1% trifluoroacetic acid was added. Extracts were passed through MultiScreen HV-HTS plates (Millipore) into a collection plate by centrifugation at 1,000 × g for 3 min and stored at 4 °C.

## LC/MS/MS

Digests were analysed at a shared instrument facility on a Paradigm MS4 nano-LC system coupled to an LCQ DECA XP ion-trap mass spectrometer (Thermo Fisher Scientific), with chromatographic separation essentially as described [2].

#### Database searching

Spectra were searched with Mascot (Matrix Science). The settings of the searches of the recombinant reference spots of Table 1, run in 2010, were: database NCBInr 20101020 (12,061,831 sequences); taxonomy *Homo sapiens*; enzyme trypsin; maximum 3 missed cleavages; mass values average; peptide mass tolerance ±1.2 Da; fragment mass tolerance ±0.6 Da; variable modification carbamidomethylation of cysteine, with phosphorylation of serine, threonine and tyrosine additionally included for one search of spot 21; no fixed modifications; protein mass unrestricted. Significance is Mascot's: individual ion scores above 43 indicate identity or extensive homology at $p < 0.05$, the threshold being 49 for the spot 21 search, which allowed the two further modifications, and Mascot applies its own identity and homology thresholds per query within a search. The full output of these searches is in Table S1.

Two further sets of searches, of spots excised from gels of endogenous 293F eluates, were run in 2014 and gave no MR1 identification; they are reported in Data Limitations. They used the same enzyme, missed-cleavage, tolerance and average-mass settings as above. The first set (gels of 2014-06, searched 2014-06) was run against NCBInr 20140323 under the taxonomies *Homo sapiens*, Mammalia, other Mammalia and Bacteria, with carbamidomethylation of cysteine as a variable modification. The second set (gels of 2014-07, searched 2014-08) was run against NCBInr 20140323 (*Homo sapiens*, Mammalia, Metazoa), NCBInr 20140906 (Mammalia, Metazoa) and SwissProt 2014_07 (Mammalia, Metazoa), with carbamidomethylation of cysteine as a fixed modification.

#### RT-PCR

RT-PCR used commercial HEK293 cDNA from the Human Cell Line MTC panel (Clontech) as template, with the primers and cycling profile described previously [1]: a single pair spanning the untranslated regions, which yields 1,083 bp for MR1A and 807 bp for MR1B, with GAPDH (983 bp) as control, PrimeSTAR HS DNA polymerase (Takara Bio) and cycles of 98 °C for 10 s, 55 °C for 15 s and 72 °C for 1 min. For this cell-line cDNA, 40 cycles were run for MR1 and 22 for GAPDH; the published protocol, which was applied to tissue cDNA panels, used 30 cycles for MR1. These results are reported in the text as data not shown.

**Table S1. Mascot output and search settings for the two-dimensional gel spots identified as MR1.** Spots were excised from the SYPRO Ruby-stained gels of Figure 2c, d and digested in-gel (Supplementary Methods). Spot numbers are the sequential positions of the excision pick list. Spots 9, 18, 21 and 24 are the spots of Table 1; spot 8 comes from a separately prepared 293-hMR1B gel that is not shown, and spot 39 lay below the rMR1B reference position and was excluded from the positional comparison. Every search matched the same database entry, `gi|3859895` (MHC class I-related protein MR1 precursor [*Homo sapiens*], nominal $M_r$ 38,586), which is the full-length MR1A sequence; the identification therefore does not distinguish the isoforms. (a) One row per search. Matches and sequences are Mascot's counts with the number significant at $p < 0.05$ in parentheses. The identity threshold is the ion score above which Mascot reports identity or extensive homology at $p < 0.05$ for that search; it is higher for the spot 21 search because that search allowed two further variable modifications. Common to all searches: MS/MS ion search; enzyme trypsin; maximum 3 missed cleavages; average masses; peptide mass tolerance ±1.2 Da; fragment mass tolerance ±0.6 Da; protein mass unrestricted; instrument type ESI-TRAP; database NCBInr 20101020 (12,061,831 sequences); taxonomy *Homo sapiens* (234,621 sequences). CAM, carbamidomethylation. No fixed modifications were set. Coverage is the sequence coverage of the Protein View report, which is available for the searches of spots 9 and 21. (b) Every peptide listed for the MR1 hit, in query order. Residues are positions in the canonical precursor of UniProt Q95460; all of them lie outside the $\alpha$3 segment (residues 203–294) deleted in MR1B, so all are shared between MR1A and MR1B. Flanking residues are shown as Mascot prints them. All peptides have no missed cleavage. Δ is $M_r$(expt) − $M_r$(calc). Significance is per query against that query's own identity and homology thresholds; "identity" and "homology" are the two significant classes at $p < 0.05$ and "–" is not significant. (c) The searches of (a) are those in which MR1 was the identification reported in the 2010 summary of all 50 excised spots, which lists the top-ranked hit for each spot. MR1 also appears as a lower-ranked protein family in two further spots, listed here; neither lies at the reference coordinates compared in the text.

(a) Searches

| Spot | Cell line | Figure | Protein score | Matches | Sequences | Identity threshold | Queries | Variable modifications | Search date | Coverage |
|---|---|---|---|---|---|---|---|---|---|---|
| 9 | 293-hMR1A | 2c | 91 | 2 (2) | 2 (2) | 43 | 839 | CAM | 26 Oct 2010 | 8% |
| 18 | 293-hMR1B | 2d | 107 | 6 (4) | 5 (3) | 43 | 830 | CAM | 26 Oct 2010 | not saved |
| 21 | 293-hMR1B | 2d | 95 | 5 (3) | 5 (3) | 49 | 852 | CAM, Phospho (ST), (Y) | 29 Oct 2010 | 22% |
| 24 | 293-hMR1B | 2d | 46 | 2 (1) | 2 (1) | 43 | 813 | CAM | 29 Oct 2010 | not saved |
| 8 | 293-hMR1B | not shown | 48 | 2 (1) | 2 (1) | 43 | 805 | CAM | 26 Oct 2010 | not saved |
| 39 | 293-hMR1B | 2d | 35 | 1 (1) | 1 (1) | 43 | 717 | CAM | 29 Oct 2010 | not saved |

(b) Matched peptides

| Spot | Query | Peptide | Residues | Observed *m/z* | $M_r$(expt) | $M_r$(calc) | Δ (Da) | Ion score | Expect | Significance |
|---|---|---|---|---|---|---|---|---|---|---|
| 9 | 419 | `R.EQNGAIYLPTPDR.-` | 329–341 | 738.0870 | 1474.1593 | 1473.5861 | 0.5732 | 64 | 2.1e-05 | identity |
| 9 | 527 | `K.DTLSWLAVDNVAHTIK.Q` | 147–162 | 893.0323 | 1784.0499 | 1782.9900 | 1.0599 | 55 | 7.9e-05 | identity |
| 18 | 78 | `R.FLEYGK.D` | 190–195 | 757.2138 | 756.2064 | 755.8577 | 0.3487 | 28 | 0.46 | – |
| 18 | 420 | `R.EQNGAIYLPTPDR.-` | 329–341 | 737.8030 | 1473.5913 | 1473.5861 | 0.0052 | 54 | 0.0041 | identity |
| 18 | 510 | `K.QAWEANQHELLYQK.N` | 163–176 | 880.0664 | 1758.1180 | 1757.8992 | 0.2188 | 7 | 1.1 | – |
| 18 | 520 | `K.DTLSWLAVDNVAHTIK.Q` | 147–162 | 892.7866 | 1783.5585 | 1782.9900 | 0.5684 | 66 | 1.7e-05 | identity |
| 18 | 789 | `R.LGVSDPIHGVPEFISVGYVDSHPITTYDSVTR.Q` | 32–63 | 1154.2830 | 3459.8267 | 3458.7825 | 1.0442 | 29 | 0.02 | homology |
| 21 | 94 | `R.FLEYGK.D` | 190–195 | 756.7087 | 755.7013 | 755.8577 | −0.1563 | 20 | 0.67 | – |
| 21 | 112 | `R.YTQLLR.G` | 84–89 | 794.1933 | 793.1859 | 792.9225 | 0.2634 | 28 | 0.52 | – |
| 21 | 538 | `K.QAWEANQHELLYQK.N` | 163–176 | 880.4841 | 1758.9535 | 1757.8992 | 1.0543 | 53 | 0.00013 | identity |
| 21 | 547 | `K.DTLSWLAVDNVAHTIK.Q` | 147–162 | 892.9707 | 1783.9266 | 1782.9900 | 0.9366 | 58 | 0.00021 | identity |
| 21 | 811 | `R.LGVSDPIHGVPEFISVGYVDSHPITTYDSVTR.Q` | 32–63 | 1154.1511 | 3459.4312 | 3458.7825 | 0.6487 | 40 | 0.0013 | homology |
| 24 | 149 | `R.YTQLLR.G` | 84–89 | 794.0813 | 793.0739 | 792.9225 | 0.1514 | 9 | 1.1 | – |
| 24 | 555 | `K.DTLSWLAVDNVAHTIK.Q` | 147–162 | 892.6392 | 1783.2635 | 1782.9900 | 0.2735 | 46 | 0.0011 | identity |
| 8 | 123 | `R.FLEYGK.D` | 190–195 | 756.9971 | 755.9897 | 755.8577 | 0.1320 | 22 | 0.88 | – |
| 8 | 550 | `K.DTLSWLAVDNVAHTIK.Q` | 147–162 | 892.6621 | 1783.3094 | 1782.9900 | 0.3194 | 48 | 0.00034 | identity |
| 39 | 112 | `R.FLEYGK.D` | 190–195 | 757.2156 | 756.2082 | 755.8577 | 0.3505 | 35 | 0.025 | homology |

(c) Spots in which MR1 appeared only as a lower-ranked family hit

| Spot | Pick-list coordinates (x, y) | MR1 family score | Identification reported in the 2010 summary |
|---|---|---|---|
| 10 | 1383, 703 | 33 | none |
| 27 | 2071, 1547 | 46 | dnaJ homolog subfamily B member 11 precursor (score 89) |

**Table S2. In silico tryptic peptides of MR1B that span the splice junction.** Peptides were enumerated from the MR1B sequence (isoform 3 of UniProt Q95460, entry version 205 of 2026-09-02: canonical precursor with the signal peptide, residues 1–22, removed and the α3 segment, residues 203–294, deleted), assuming cleavage C-terminal to Lys and Arg but not before Pro, and up to three missed cleavages (MC), the maximum allowed in the searches of Table S1. Length is the number of residues. Masses are of the unmodified neutral peptide (sum of residue masses plus the mass of water); they are neither protonated masses nor m/z values, and no modification, for example oxidised Met, is included. Group A comprises the peptides that terminate at Lys303 and do not extend into the following hydrophobic stretch (residues 304–323); group B those that extend past Lys303 and therefore include the entire annotated transmembrane region (residues 303–323), there being no Lys or Arg within residues 304–323. Residue numbers refer to the canonical precursor. The last column states whether the average mass falls within 700–4000 Da; this window was adopted for the present theoretical comparison and is not the acquisition range of the instrument used, nor does membership in it indicate that a peptide was detectable. The two peptides marked as outside exceed 4000 Da in both mass conventions. None of these peptides was reported in the searches of Table S1; the factors that determine whether they would have been observed were not measured here (see Data Limitations and Perspectives in the main text). For the first candidate of group A, `TESETIPLVMK`, the doubly protonated ion is at m/z 624.33 (monoisotopic).

| Peptide | MC | Length | Average mass (Da) | Monoisotopic mass (Da) | 700–4000 Da |
|---|---|---|---|---|---|
| **Group A: terminating at Lys303** | | | | | |
| `TESETIPLVMK` | 0 | 11 | 1247.47 | 1246.65 | within |
| `DTLQRTESETIPLVMK` | 1 | 16 | 1861.14 | 1859.97 | within |
| `FLEYGKDTLQRTESETIPLVMK` | 2 | 22 | 2598.99 | 2597.34 | within |
| `RFLEYGKDTLQRTESETIPLVMK` | 3 | 23 | 2755.18 | 2753.44 | within |
| **Group B: including residues 303–323** | | | | | |
| `TESETIPLVMKAVSGSIVLVIVLAGVGVLVWR` | 1 | 32 | 3337.07 | 3334.93 | within |
| `TESETIPLVMKAVSGSIVLVIVLAGVGVLVWRR` | 2 | 33 | 3493.25 | 3491.03 | within |
| `TESETIPLVMKAVSGSIVLVIVLAGVGVLVWRRRPR` | 3 | 36 | 3902.74 | 3900.29 | within |
| `DTLQRTESETIPLVMKAVSGSIVLVIVLAGVGVLVWR` | 2 | 37 | 3950.74 | 3948.25 | within |
| `DTLQRTESETIPLVMKAVSGSIVLVIVLAGVGVLVWRR` | 3 | 38 | 4106.92 | 4104.35 | outside |
| `FLEYGKDTLQRTESETIPLVMKAVSGSIVLVIVLAGVGVLVWR` | 3 | 43 | 4688.59 | 4685.62 | outside |

The four peptides of group A are nested sequences that share the splice junction, Met302 and the C-terminal Lys303, and differ only in missed cleavages on their N-terminal side. They are specific to MR1B among the annotated isoforms of UniProt Q95460.

**Figure S1. Uncropped membrane and exposure series for Figure 1.** The one-dimensional blot of 2014-05-14, imaged on an ImageQuant LAS 4000 at the four exposures recorded and as the 400 s cumulative image. The membrane carried twelve loadings: the flow-through of the concentration step, the MagicMark XP marker, one blank, and three groups of three sample lanes (293-hMR1A lysate, 293F immunoprecipitation eluate, 293-hMR1B lysate) probed with NB0546, with NB0560, and with NB0560 pre-incubated with the competing peptide. The membrane was cut between the groups so that each piece could be probed separately and the pieces were imaged together, which is why the groups are separated by blank strips. Figure 1 shows the nine sample lanes; the flow-through and marker lanes were cropped from it. Each panel is one whole frame, with no part cut out, moved, inserted or spliced, and the first five have had no level, brightness, contrast or gamma adjustment: the 16-bit originals are rendered to 8-bit linearly. The last panel of each series repeats the 400 s cumulative image with a linear contrast stretch applied uniformly to the whole frame, the brightest pixel mapped to white and a value 9,600 counts below it to black on the 16-bit scale that runs to 65,535, with no gamma and nothing local. The stretch saturates the inside of the strongest spots, and the unadjusted panel of the same exposure is shown beside it. Panels are shown in the orientation in which the instrument displays the membrane, which is the orientation of the published figures; the published panels carry their own brightness and contrast settings and so look stronger than the unadjusted panels here.

10 s | 30 s | 90 s | 270 s | 400 s (cumulative) | 400 s, contrast stretched

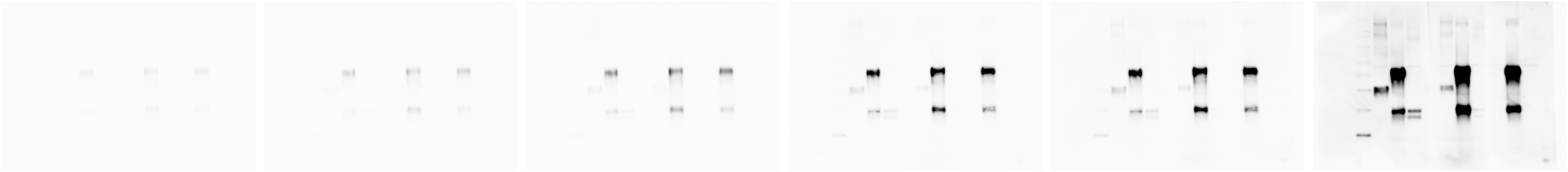

**Figure S2. Uncropped images for Figure 2.** (a) The two-dimensional blot of 2014-05-21, imaged at the four exposures recorded and as the 400 s cumulative image. Four membranes were imaged together in one frame: the 293-hMR1A eluate (upper left) and the 293-hMR1B eluate (upper right), which are Figure 2a and Figure 2b, and the parental 293F eluate and a flow-through control (lower left and lower right), which are not shown in the paper. Each panel is one whole frame, with no part cut out, moved, inserted or spliced. Panels are treated as in Figure S1. (b) is on the next page.

(a)

10 s | 30 s | 90 s | 270 s | 400 s (cumulative) | 400 s, contrast stretched

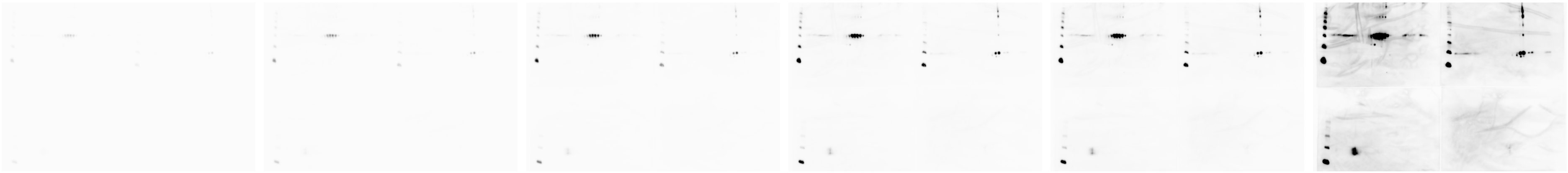

**Figure S2.** **(continued)** (b) The SYPRO Ruby fluorescence scan of 2010-10-19 from which Figure 2c and Figure 2d were cropped, shown whole and in the same orientation as those panels. The six gels of the two runs were laid out together in two columns of three, fixed at the edges with agarose, provided with the two reference markers visible as dots at the top and bottom edges, and scanned on a Typhoon imager at a 100 µm pixel size before spot excision. In this orientation the handwritten labels of the original read sideways, so the gels are identified here by position, which was checked against the coordinates of the excision pick list: Figure 2c is the middle gel of the right column, 101006 hMR1(full), and Figure 2d the bottom gel of the left column, 101016 hMR1(a3−); Figure 2c was additionally rotated by approximately 1.8° for the paper. The two parental 293F control gels referred to in the text as showing no spots at the reference positions are the top gels of the left column (101016) and the right column (101006), and are shown here in full; the remaining two are 101016 hMR1(full), middle left, and 101006 hMR1(a3−), bottom right. This is a single fluorescence scan of all six gels, shown whole, so that the relative position of every gel and every spot can be checked against the pick list, and with no level, brightness, contrast or gamma adjustment.

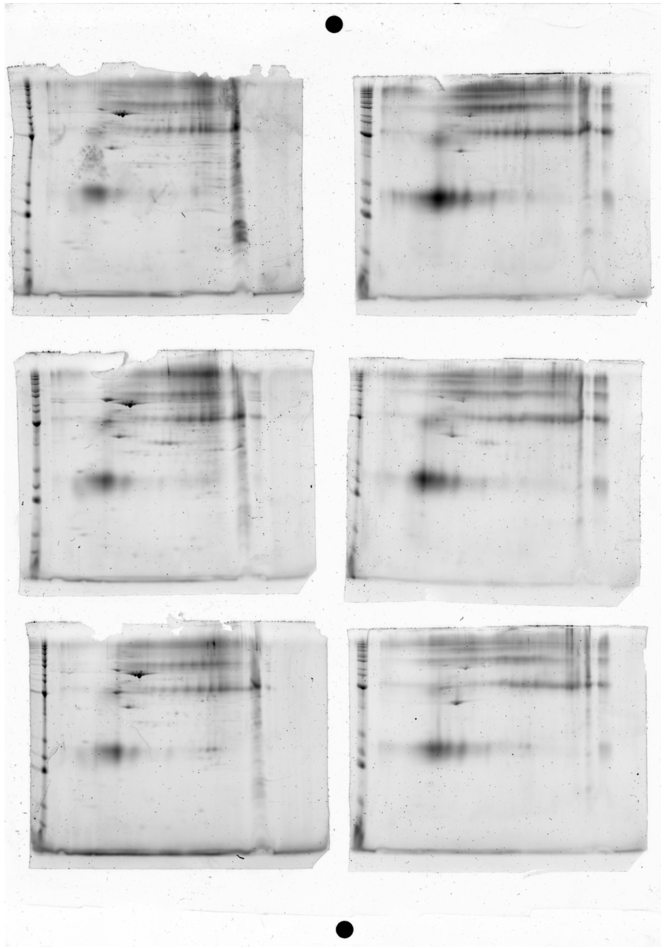

**Figure S3. Uncropped membranes and exposure series for Figure 3.** The three two-dimensional blots of the Figure 3 preparation, each imaged at the four exposures recorded and as the 400 s cumulative image. (a) 2014-11-13, the two pre-immune control membranes, which were imaged together in one frame and are shown here in that one frame: the upper membrane is Figure 3a and the lower, carrying the handwritten label 141113 NB0560 pre, is Figure 3b. The strong train of spots on each is the immunoglobulin-related signal referred to in the text, and it is present in the published panels as well. (b) and (c) are on the next page. Every panel is one whole frame, with no part cut out, moved, inserted or spliced, so each shows every mark on the membrane at that exposure. Panels are treated as in Figure S1. The frame of (a) is turned a quarter turn into the orientation of the published panels; no other geometric change was made.

(a) 2014-11-13, pre-immune sera (Figure 3a, b)

10 s | 30 s | 90 s | 270 s | 400 s (cumulative) | 400 s, contrast stretched

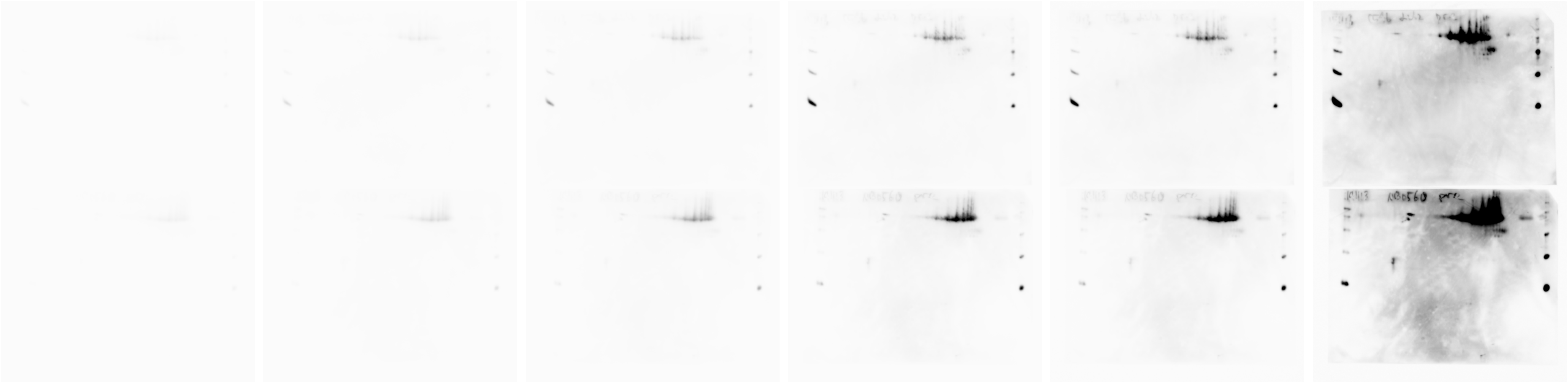

**Figure S3. (continued)** (b) 2014-08-15, NB0546: Figure 3c is a crop of this frame. (c) 2014-11-12, NB0560: Figure 3d is a crop of this frame. Conditions and treatment as in (a).

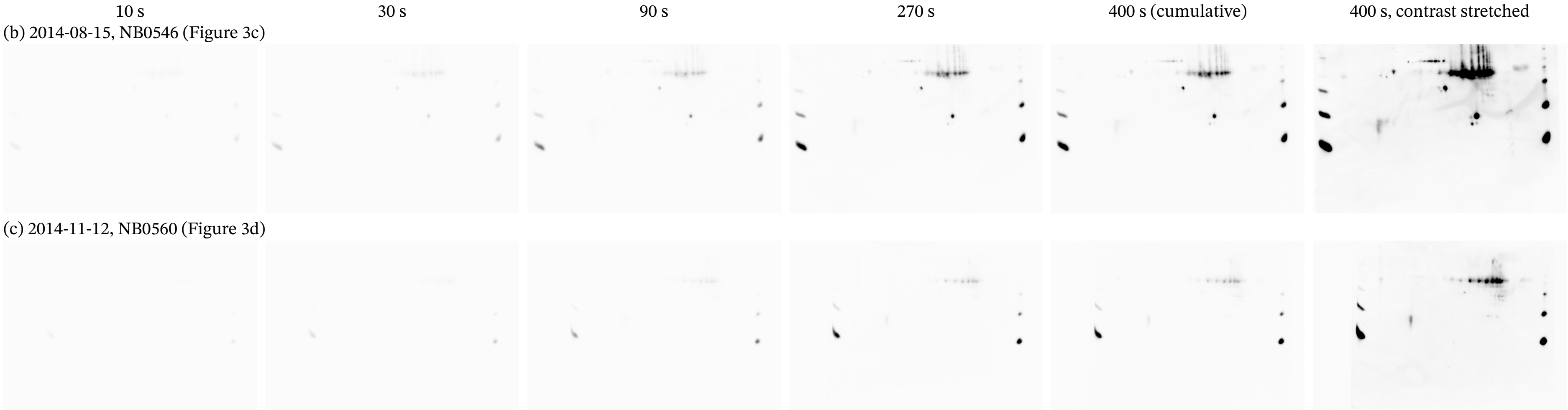

### References

[1] H. Yamaguchi, K. Tsukamoto, K. Hashimoto, Cell surface expression of MR1B, a splice variant of the MHC class I-related molecule MR1, revealed with antibodies, Biochem. Biophys. Res. Commun. 443 (2014) 422–427. https://doi.org/10.1016/j.bbrc.2013.11.096.

[2] G. Kurosawa, M. Sumitomo, Y. Akahori, et al., Methods for comprehensive identification of membrane proteins recognized by a large number of monoclonal antibodies, J. Immunol. Methods 351 (2009) 1–12. https://doi.org/10.1016/j.jim.2009.09.003.
